# bayesynergy: flexible Bayesian modelling of synergistic interaction effects in in-vitro drug combination experiments

**DOI:** 10.1101/2021.04.07.438787

**Authors:** Leiv Rønneberg, Andrea Cremaschi, Robert Hanes, Jorrit M. Enserink, Manuela Zucknick

## Abstract

The effect of cancer therapies is often tested pre-clinically via *in-vitro* experiments, where the post-treatment viability of the cancer cell population is measured through assays estimating the number of viable cells. In this way, large libraries of compounds can be tested, comparing the efficacy of each treatment. Drug interaction studies focus on the quantification of the additional effect encountered when two drugs are combined, as opposed to using the treatments separately. In the **bayesynergy** R package, we implement a probabilistic approach for the description of the drug combination experiment, where the observed dose response curve is modelled as a sum of the expected response under a zero-interaction model and an additional interaction effect (synergistic or antagonistic). The interaction is modelled in a flexible manner, using a Gaussian process formulation. Since the proposed approach is based on a statistical model, it allows the natural inclusion of replicates, handles missing data and uneven concentration grids, and provides uncertainty quantification around the results. The model is implemented in the Stan programming language providing a computationally efficient sampler, a fast approximation of the posterior through variational inference, and features parallel processing for working with large drug combination screens.

## 1 Introduction

In pre-clinical cancer drug sensitivity screening, the effectiveness of compounds is tested *in-vitro* on cell lines or samples derived from patients. The response of those cell lines or patient-derived samples to a treatment is measured with dose-response experiments, in which cells have been exposed to a range of drug concentrations over a period of time. The output is typically a measure of cell viability or other type of cell count obtained from single-drug experiments, which are used to model the response by fitting a parametric log-logistic model to the concentration-response curve, e.g

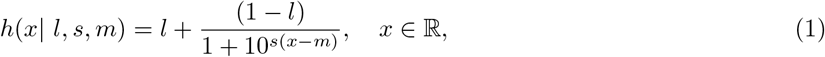

where *x* denotes the log_10_ drug concentration, as it is assumed throughout the paper. We utilize a three parameter log-logistic curve rather than the more common four parameter version, fixing the upper asymptote to one. Working with properly normalized viability measurements, this simply reflects that at very low levels of concentration, there is completely unhindered cell growth. Positive values of the slope parameter *s* are associated with measuring inhibition of cell viability. The parameter *l* controls the lower asymptote, or the maximum efficacy of the drug. Some anti-cancer drugs are cytostatic rather than cytotoxic, leading to a plateau effect in the dose-response curve. The lower asymptote captures this potential diminishing return of increased drug concentration. The inflection point of the curve, *m*, corresponds to the concentration of the compound needed to induce a response equal to 50% of the maximum response and is commonly referred to as *half-maximal effective concentration* (or EC_50_), a popular measure of efficacy of the compounds.

In drug combination studies, more than one compound is tested at the same time, with the aim of finding more effective treatments. In particular, one is interested in identifying drug combinations that are either *synergistic* or *antagonistic*. An interaction between two drugs resulting in a combined effect greater than expected is called synergistic, while an effect lower than expected is called antagonistic. The expected combined effect of two drugs hinges on an assumption of how the two drugs might behave jointly, having only been observed individually. Several such assumptions exist, each having its own underlying pharmacological reasoning. Starting from these assumptions the output of drug combination experiments can be modelled using suitable mathematical models with the aim of quantifying the interaction component. Building on work dating back to the first half of the twentieth century [see for instance 1, 2], the literature on the topic has developed widely in recent years [see 3, 4, 5, for reviews on the topic].

Several software packages for analysing drug sensitivity data exist, notably the **R** packages **synergyfinder** for drug combination studies [6, 7], and **drc** for single drug data [8]. In addition, standalone software such as **Combenefit** [9] has been utilised to quantify interaction effects in high-throughput drug combination experiments, where thousands of experiments are analysed simultaneously.

A common drawback of all classical drug interaction models implemented in these packages is that they interpret any deviation of the observed data from the expected non-interaction model as interaction, i.e. as evidence for synergistic or antagonistic effects. Therefore, these models do not allow for heterogeneity in the data, for measurement errors or for any other biological or technical variation, which are commonplace in high-throughput data. A notable exception is the work of [10], where a framework accounting for experimental noise is developed, but it only models the experiments point-wise, not considering the full dose-response surface.

To overcome these problems, the **R** package **bayesynergy** implements a statistical model for studying the interaction between two drugs, where the drug combination surface is modelled using a flexible Bayesian approach. This formulation allows for proper inclusion of data variability, uses replicate measurements when available, and naturally handles missing data and uneven concentration grids. The **bayesynergy** package implements an extension of the model developed in [11], where the drug response surface is interpreted as the result of a stochastic model, able to discriminate between its zero-interaction and interaction parts. While the zero-interaction part is given a parametric model, corresponding to the product of the dose-response curves estimated for each drug individually, the interaction part is modelled in a nonparametric fashion using a Gaussian Process.

## 2 Model

The main focus in drug-response studies is the dose-response function, *f* (*x*), that maps drug concentration *x* to a measure of cell survival, e.g. the percentage of cells still viable after treatment. Utilizing the percentage of survival, it is assumed that *f* only takes values in the interval (0, 1), the boundary reflecting complete cell death, or complete cell survival.

From this function, numerous summary measures can be derived, for example the half maximal inhibitory concentration (EC_50_), or the drug sensitivity score (DSS) [12], both attempting to quantify a compound’s efficacy. In drug combination studies, ***x*** = (*x*_1_, *x*_2_) denotes the pair of concentrations of the two drugs being combined, and it is assumed that the drug response function can be decomposed as:

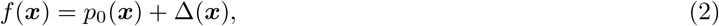

where *p*_0_(***x***) is the non-interaction effect, and Δ(***x***) is the interaction effect. While the drug response function cannot be directly observed, experimenters can obtain noisy evaluations of it through e.g. cell viability assays.

The non-interaction effect *p*_0_ encodes an assumption on how the drugs would behave together, if they truly did not interact. For this term, we assume a Bliss independence model [2], which corresponds to a probabilistic independence assumption on the joint effect. That is, if we interpret the log-logistic curves from equation (1) as probabilities of a cell’s viability at concentration *x* of a drug, the joint probability of a cell’s ability to proliferate at concentration ***x*** takes the following form:

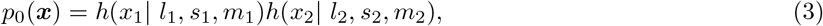

where *l*_*i*_, *s*_*i*_, *m*_*i*_ for *i* = 1, 2 are the individual parameters for the two drugs being combined, introduced in (1).

The interaction effect Δ captures any additional effect of the two drugs in combination that is not captured under the non-interaction assumption by *p*_0_. In order to ensure the flexibility of the drug response function this term is given a zero-mean Gaussian Process (GP) prior [13]. A GP is a stochastic process, any finite realization of which is distributed as a multivariate normal. That is, if

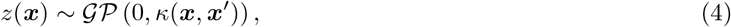

then

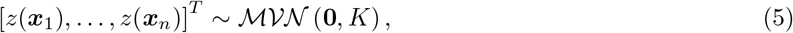

where the entries of the covariance matrix are *K*_*ij*_ = *κ*(***x***_*i*_, ***x***_*j*_). The function *κ*(·,·) is called the kernel, or covariance function, and encodes the smoothness on the final function. We will assume a stationary kernel that only depends on the distance between covariates, such that:

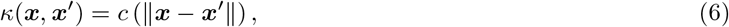

for some function *c*(*·*).

Furthermore, to ensure that the resulting dose-response function only takes values within the interval (0, 1), the Gaussian process is bounded via a transformation function:

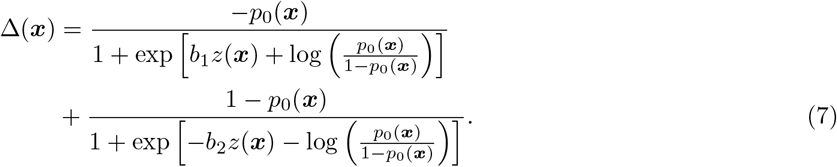

The parameters (*b*_1_, *b*_2_) are not directly interpretable, but help keep the model identifiable by imposing *b*_1_ ≠ *b*_2_ through separate continuous prior distributions.

In addition to giving the correct bounds for the dose-response function, the transformation function acts to provide a slightly conservative prior distribution on Δ(**x**). The underlying GP has mean zero so that substituting *z*(**x**) = 0 in the equation above yields Δ(*x*) = 0, ensuring that as the underlying GP reverts to its prior in absence of data, the dose-response function will revert to the non-interaction assumption. This reflects our belief that true interaction is a rare occurrence, and we build the model with minimal bias towards it. However, due to the non-linear transformation function, in terms of prior expectation we only have 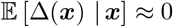. See the appendix for more details.

### 2.1 Observation model

Cell viability is typically measured *in-vitro* using various cellular assays. In these assays, viability is determined indirectly by measuring a marker associated with cell viability (e.g. ATP levels) and comparing these to measurements taken from negative and positive controls. Once properly normalized, these measurements can be thought of as evaluations of the underlying dose-response function *f*. However, due to technical and biological noise sources, it is not uncommon to get viability measurements outside of the interval (0, 1). To minimize the influence of noise on the output, experiments are frequently performed including replicate observations.

In order to take this into account we assume:

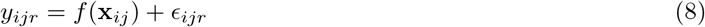

where ***x***_*ij*_ = (*x*_1*i*_, *x*_2*j*_) denotes the concentration pairs used in the drug combinations, where measurements are indexed by *i* = 1, … , *n*_1_ and *j* = 1, … , *n*_2_, while the subscript *r* = 1, … , *n*_rep_ denotes the replicates. The noise term *ϵ*_*ijr*_ captures the measurement error around the dose-response curve, and is given a zero-mean normal distribution, independent across replicates, with a heteroscedastic variance:

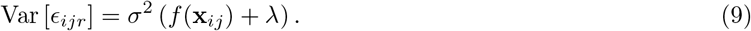

A similar structure was used in [14] directly, from a log-transformation, and in [15] indirectly, through modelling the raw fluorescent intensity output from the plate reader. The heteroscedastic structure arises from the normalization procedure itself. Positive controls typically have much lower variance than the negative controls, i.e. it is much easier to establish when most of the cells are dead, rather than alive.

In the positive controls, a cytotoxic compound has been added to ensure complete cell death. The variance across measurements from these controls reflect a baseline level of technical noise, i.e. measurement noise introduced by the instrument itself. In the negative controls cells are allowed to proliferate freely, not experiencing inhibitory effects from chemical compounds. The variation across the negative controls reflects the heterogeneity of cell growth, in addition to the underlying level of technical noise. Coupling these controls to the dose-response function *f*, the positive controls reflect the case where all the cells are dead, i.e. *f* = 0, while the negative controls provide the case where all cells are still alive, i.e. *f* = 1.

Letting the dose-response function vary from *f* (**x**) = 1, the setting of the negative controls, to *f* (**x**) = 0, the setting of positive controls, *σ*^2^ can be thought of as the overall biological heterogeneity, while *λ* is added as an offset, for the technical noise. We add *λ* inside the parenthesis, to be multiplied by *σ*^2^ as it is more robust against model misspecification. The parameter *λ* is not given a prior distribution, but must be set by the user to reflect the data at hand. Letting 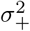 and 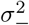 denote the variances of the positive and negative controls respectively, the value of *λ* can be set empirically as:

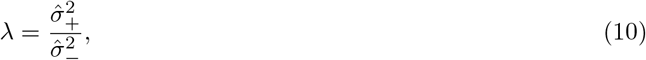

where 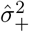 and 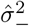 are estimates obtained from the positive and negative controls. Ideally, we would like to set *λ* equal to the ratio between the technical noise and biological heterogeneity, but we do not have access to the biological heterogeneity directly. These quantities should be familiar to experimenters working with these data, as they are key ingredients in calculating quality control measures of the assay, such as the Z-prime factor [16].

A full model specification, with all prior distributions can be found in the appendix. The prior choices are inherently linked to the drug concentration ranges we see in cancer drug screens. Concentrations are typically given in micro-molars (*μ*M), and are equally spaced out on the log_10_ scale. A typical range of concentrations in a large drug combination screen would be from as low as 10^−6^*μ*M (picomolar) to as high as 10^3^*μ*M (millimolar), for which the prior distributions should be sufficiently calibrated.

## 3 Implementation

The model is implemented using the R interface of the Stan programming language [17, 18]. Posterior samples are obtained using Markov Chain Monte Carlo (MCMC), specifically a version of Hamiltonian Monte Carlo called the ‘No U-Turn Sampler’ (NUTS) [19]. In addition, Stan provides an algorithm for variational inference called Automatic Differentiation Variational Inference (ADVI) [20], that provides a quick approximation to the posterior distribution.

Gaussian Processes can be computationally expensive in a fully Bayesian setting, with *n* data points requiring the Cholesky decomposition of an *n × n* matrix at each step of the sampling scheme. Thanks to the grid structure in drug combination experiments, we can significantly speed things up. Following the Stan implementation given in [21], we start by writing the kernel function of the GP as:

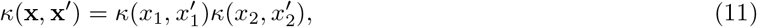

where 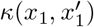 and 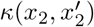 denotes kernel functions defined on the pairwise individual drug concentrations. From this structure the covariance matrix *K* from equation (5) can be written as a Kronecker product, 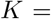 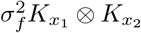, where 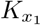 is an *n*_1_ × *n*_1_ matrix with entries 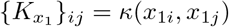, and similarly for the *n*_2_ × *n*_2_ matrix 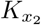.

By utilizing the following property:

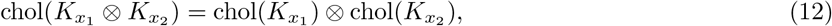

the calculation of the GP only requires the Cholesky decomposition of the two smaller matrices 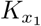 and 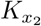 of dimension *n*_1_ × *n*_1_ and *n*_2_ × *n*_2_, respectively. Samples from the GP are obtained using a latent formulation, where we first create an *n*_2_ × *n*_1_ matrix *V* whose entries are standard normal latent variables, and create the matrix *Z* as:

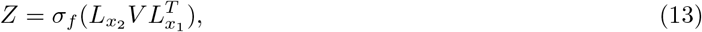

where 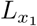 and 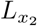 are the Cholesky decompositions of 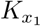 and 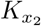, respectively. This ensures that vec(*Z*), where vec() denotes the operator that creates a column vector by stacking the columns together, has the required multivariate normal distribution with the specified Kronecker covariance structure.

### 3.1 Missing data

The fast speed-ups offered by the covariance structure requires a full matrix of latent factors *V*_*ij*_, one for each combination of drug concentrations, but crucially it does not require an observed viability score at each location.

In real datasets, there are many reasons why we might not have access to a full grid of viability scores, or an equal number of replicates at each location. Most commonly, resource constraints prohibit a full exploration of the dose-combination landscape, some researchers opting for sparse designs, where only some of the concentration combinations are actually observed.

Some typical designs are visualised in Figure 1. In the first panel (a) we see the “full” design, where every combination of mono-therapy concentration has also been observed for the combinations. This is the ideal setting that very often is not achieved in real datasets. The next two panels (b and c) show the designs where one or both drugs are fixed at a single concentration – we call these the “line” or “cross” design, respectively. These have been employed successfully in experiments using patient derived samples where the number of cells available are limited, e.g. on leukaemia [22]. The next panel (d) denotes the “diagonal” design, promoted in [23], who propose a machine-learning algorithm for imputing the full matrix.

**Figure 1:**
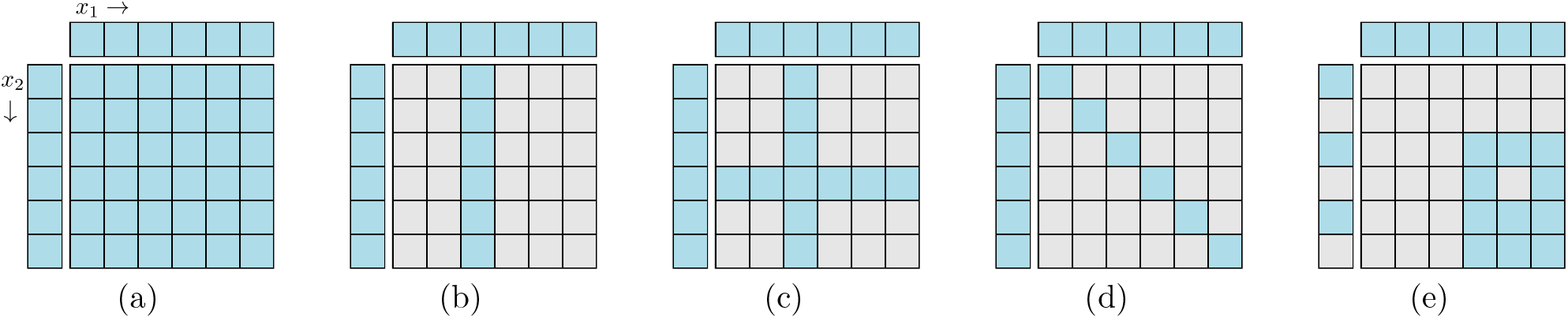
Various experimental designs common in drug-combination experiments. The disconnected first row and column indicates values of the mono-therapies, with concentrations increasing along the rows and columns as indicated in panel (a). The inside of the matrix corresponds to the combination measurements, where a blue colour indicates that the viability has been observed for this location, whilst grey indicates combinations that are missing. Various patterns of missingness give rise to a fully observed (a), line (b), cross (c), diagonal (d), or shifted design (e).

The last panel (e) shows the situation where the mono-therapy experiments have been performed at a different concentration grid than the combination experiment – we denote this the “shifted” design. This is the design utilised in [24], one of the publicly available datasets for drug combinations. The panel also shows a common situation where a single observation has been removed, possibly after being deemed an outlier. Finally, there is a concentration of drug 2 (a row) that has no observations at all, neither for the mono-therapy nor for the combination. Essentially a prediction task, this is simply handled as another instance of missing data. This is useful, if one wishes to compute the dose-response function on a finer grid of concentrations, or outside the range of the data.

The implementation supports any pattern of missing data, and provides posterior estimates of the dose-response function *f*, evaluated at every combination of concentrations ***x*** it is given. Missing entries are imputed through the sampling process and provided with full uncertainty quantification.

### 3.2 Large drug combination screens

In the setting of large drug combination screens, where thousands of experiments need to be pushed through an analysis pipeline, computational speed can quickly become a significant bottleneck. The bayesynergy package contains functionality for parallel processing in the setting of large screens, and provides automatic error checking and retries in the case of poor model fits. The user is given various flags to indicate whether an experiment needs closer inspection, e.g. to correct an error in the input format, and failed experiments can be easily fed back into the pipeline again. See the bayesynergy package vignette for more details on how to diagnose warnings and error messages.

The computational time of a single experiment depends on the number of unique combination of concentrations, the pattern of missing data, and the number of replicates. Though the Kronecker-structured covariance matrix speeds things up significantly, the computation time of individual experiments can be further improved by using variational inference. The ADVI algorithm provides an approximation to the full posterior distribution, and gives a rough estimate of the model parameters, often orders of magnitude faster than full posterior sampling via the NUTS algorithm. In our experience, the variational approximation is relatively accurate as an initial exploration of large datasets, and it is able to identify interesting experiments, e.g. those with large synergistic regions. These experiments can then be followed up by running the more expensive NUTS algorithm for full posterior sampling.

## 4 Posterior summaries

Given an input of viability scores and drug concentrations, the `bayesynergy` package provides inference for the joint posterior distribution of all model parameters, either by generating samples from the posterior distribution using the NUTS sampler, or by approximate inference via the variational inference algorithm ADVI. From these, we construct samples from the posterior dose-response function *f*, and its constituent parts *p*_0_ and Δ, evaluated at every combination of the drug concentrations given as input (see Figure 2). From these matrices, further summary measures of drug response can be quantified.

**Figure 2:**
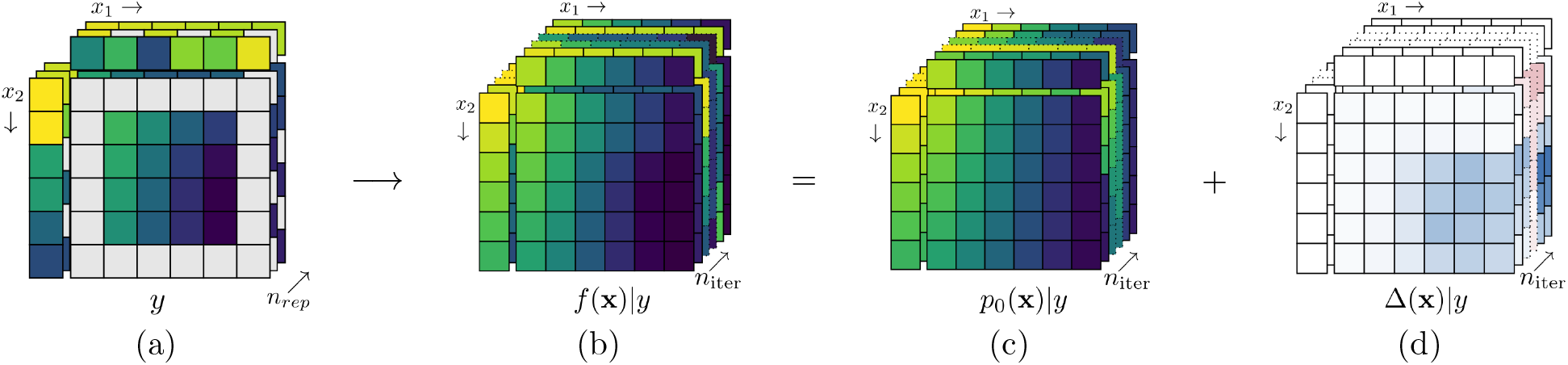
Given an input of *n*_rep_ incomplete dose-response matrices (a), perhaps with different patterns of missingness, the bayesynergy function returns *n*_iter_ samples from the posterior dose-response function *f* (b), split into its non-interaction (c) and interaction (d) parts, each evaluated on the complete set of inputs.

Summarising the drug response into a single number is useful when comparing different treatment options, or as input to other algorithms, e.g. for prediction purposes. Because of the measurement error inherent in cell viability screens, these summaries are themselves quite noisy, and care must be taken when comparing values across cell samples. The uncertainty in summary statistics can be gathered using e.g. 95% credible intervals (CI), to better discern true synergistic effects from background noise.

From the posterior samples of the dose-response function we produce a number of summary measures, each with corresponding uncertainty. From the mono-therapy curves, in addition to the EC_50_ parameter *m*, we compute DSS scores, that are normalized to the drug concentration range. In addition, we produce various efficacy measures for the drug combination, including a two-dimensional version of the DSS, called the residual Volume Under Surface (rVUS), introduced in [11]. The general equation for rVUS is defined by the double integral:

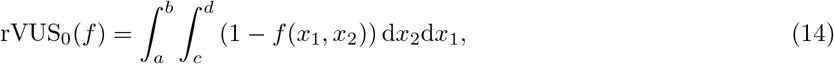

and gives a measure of the total efficacy of the two drugs combined. The integration limits are given by the minimum and maximum drug concentrations, i.e. *a* = min (*x*_1_), *b* = max (*x*_1_), *c* = min (*x*_2_) and *d* = max (*x*_2_). The value is further normalized by:

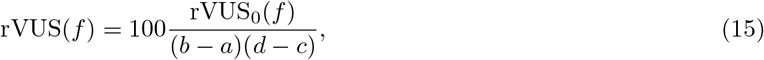

and becomes interpretable as the percentage of a hypothetical *maximum efficacy* that the drug combination achieved over the combined concentration range. Similarly, we compute this efficacy measure for both the non-interaction surface and the interaction surface, splitting the efficacy into its two main contributions. Because the interaction surface can be complex, with local regions of synergy and antagonism that can cancel each other out when taking the integral, we further compute rVUS(Δ^−^) and rVUS(Δ^+^). Here Δ^−^ and Δ^+^ denote the synergistic and antagonistic parts of the interaction surface, i.e. the parts of Δ above or below zero, respectively. Finally, in order to make comparisons of synergistic effects across experiments, we can compute a synergy score by standardising the rVUS(Δ^−^) by its uncertainty:

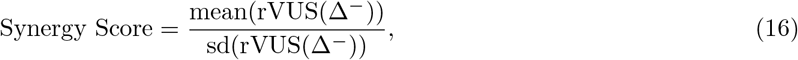

and similarly defined for an ‘Antagonism Score’. These scores can be used to rank experiments by synergy, while at the same time taking into account varying levels of uncertainty, or used as input in other algorithms.

## 5 An example workflow for a drug combination screen

To illustrate a typical analysis workflow of the `bayesynergy` package, we utilise a subset of the data provided by [24]. The full dataset contains 38 compounds screened pairwise in 583 combinations across 39 cancer cell lines. From this dataset, we select 6 breast cancer cell lines for further investigation, yielding 3 498 experiments to analyse. Each experiment usually contains 160 observations from the dose-response function, calculated on a 8 unique concentrations for the single drugs, and a 4 × 4 grid for the combinations. With the combination concentrations slightly shifted, this yields a 12 × 12 grid of possible combinations for the full grid, of which typically only 32 are actually observed, see Figure 3. Finally, these observations are made with different numbers of replicates, single drug viabilities usually replicated six times, and combination viabilities four.

**Figure 3:**
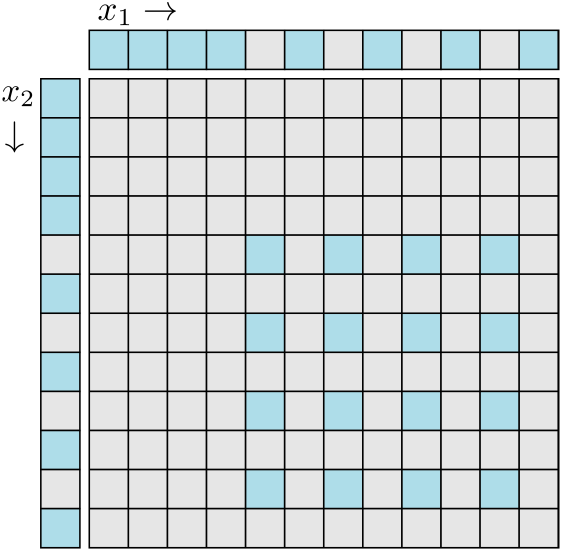
The dataset from [24] is given in a ‘shifted’ format, where the mono-therapy experiment and combination experiment has been performed on two distinct concentration grids.

Using the built-in function for analyzing large drug combination screens, `synergyscreen`, we combine parallel processing and variational approximation to fit the model to each experiment. For the 3 498 experiments, the whole process took approximately 2 hours on a 2,2 GHz Dual-Core Intel Core i7 computer, using 4 threads, at an average of 2 seconds per experiment. Being highly parallel, the computation time mainly depends on the number of cores available to the user.

This data frame can be plotted to give a quick overview of the interesting combinations. Figure 4 gives an overview of the drug combinations for the six breast cancer cell lines. The synergy and antagonism scores from equation (16) are averaged across the cell lines and colored to indicate the top synergistic or antagonistic combinations. The size of the dots indicate the median average deviation (MAD), high values of which represent drug combinations that show very selective interaction, perhaps only in a single cell line.

**Figure 4:**
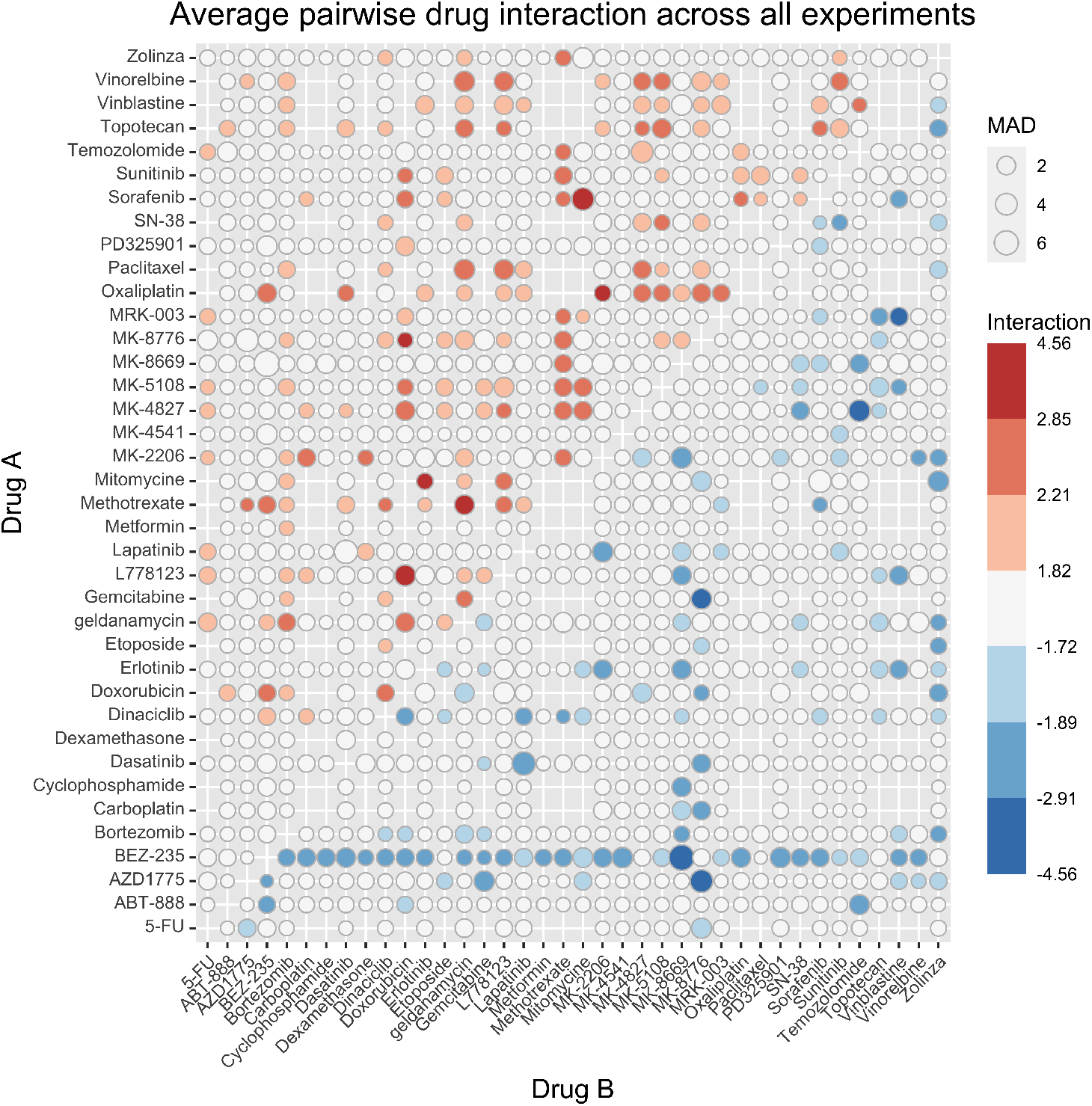
The output of `synergyscreen` can be plotted to get a quick overview of the drug screen. In this figure, synergy and antagonism scores have been averaged across the six breast cancer cell lines in [24], to produce a pairwise plot of the interaction in all combinations. The upper red triangle show antagonism scores, while the lower blue contains synergy scores. The size of the dots is proportional to the median average deviation (MAD), large values of which indicate that the combination achieves divergent scores across the cell lines.

From the plot, several combinations appear interesting for further analysis. The drugs BEZ-235 and MK-8669 are both mTOR inhibitors, with BEZ-235 being a dual mTOR/PI3K inhibitor, and produce synergistic signals across a wide range of combinations. Other drugs are more selective, only interacting with a few others.

For example, the DNA damaging agent Gemcitabine shows strong interaction only with MK-8776, a selective Chk1 inhibitor. When combined, these drugs produce a strong synergistic effect in three of the six breast cancer cell lines, which all have TP53 mutations. MK-8776 is known to enhance the rate of cell death induced by chemotherapy agents including gemcitabine [25]; the treatment combination has been tested in a phase I trial [26]. The combination of Chk1 inhibition and DNA-damaging treatment can achieve selectivity towards p53-deficient cancer cells by synthetic lethality [27].

**Table 1:** Table showing posterior estimates for parameters and summary statistics of the drug combination Gemcitabine + MK-8776 on the OCUBM cell line (data from [24]). The samples were generated using the NUTS algorithm on default model settings, with *n*_iter_ = 4000. The parameters are grouped into their corresponding location within the model hierarchy. For the user, the summary statistics at the end is the more interesting output of the model, together with perhaps the *EC*_50_ parameters, all of which is marked in bold.

| | mean | 2.5% | 97.5% | $n_{\text{eff}}$ |
| --- | --- | --- | --- | --- |
| *Observation model* |  |  |  |  |
| $\sigma$ | 0.0864 | 0.0769 | 0.0968 | 6755 |
| *Non-interaction* |  |  |  |  |
| $l_1$ (Gemcitabine) | 0.07 | 0.04 | 0.10 | 3171 |
| $l_2$ (MK-8776) | 0.64 | 0.58 | 0.69 | 3909 |
| $s_1$ (Gemcitabine) | 2.38 | 1.97 | 2.83 | 3571 |
| $s_2$ (MK-8776) | 1.94 | 1.07 | 3.37 | 4024 |
| **$m_1$ (Gemcitabine)** | -2.68 | -2.72 | -2.64 | 3238 |
| **$m_2$ (MK-8776)** | -0.53 | -0.71 | -0.33 | 3163 |
| *Interaction* |  |  |  |  |
| $\ell$ | 1.68 | 0.96 | 3.09 | 1334 |
| $\sigma_f^2$ | 3.69 | 1.32 | 13.35 | 1859 |
| *Summary statistics* |  |  |  |  |
| **DSS (Gemcitabine)** | 45.91 | 42.48 | 49.10 | 3485 |
| **DSS (MK-8776)** | 20.71 | 16.66 | 24.85 | 4419 |
| **rVUS(f)** | 53.39 | 37.18 | 72.59 | 2920 |
| **rVUS(p<sub>0</sub>)** | 29.17 | 27.47 | 30.86 | 5876 |
| **rVUS($\Delta^-$)** | 25.14 | 11.85 | 43.46 | 2816 |
| **rVUS($\Delta^+$)** | 0.93 | 0.03 | 4.47 | 3242 |

### 5.1 Detailed single experiment analysis

Inspecting this combination further, we find that it is most synergistic in the OCUBM cell line, the variational approximation reporting a point estimate of 12.28 (rVUS(Δ^−^)), and a synergy score of 5.24, indicating an effect far above the estimation uncertainty. We run the full MCMC algorithm for this experiment to obtain 4000 samples from the posterior distribution, which can be further processed and used to calculate summary statistics and produce various plots. Table 5 shows posterior means, 95% credible intervals and the effective sample size for a subset of parameters in the model in addition to the summary measures derived in the previous section, while Figure 5 shows some of the plots produced for this particular experiment. From the table, parameter estimates for the two mono-therapy curves can be examined, and the model fit can be inspected in panel (a) and (b) of the figure. With 9 replicates at each location, the mono-therapy parameters are very well estimated, with small credible intervals and high effective sample size. In the figure, the heteroscedastic nature of the observation model also becomes clearly visible. From the samples of the dose-response function as in Figure 2, summary measures of efficacy are computed and reported in Table 5. The DSS scores have rather sharp posteriors, being a bit more precise for Gemcitabine, reflecting the overall certainty in the parameter estimates. The DSS score for Gemcitabine is estimated at 45.9 with a 95% CI (42.5,49.1), while MK-8776 has a DSS score of 20.7 (16.7,24.8), indicating a lower efficacy compared to Gemcitabine. The mono-therapy curve of MK-8776 appears to plateau at around 60% viability (posterior mean of *l*_2_ at 0.64), which could indicate that the drug has saturated its target (Chk1).

**Figure 5:**
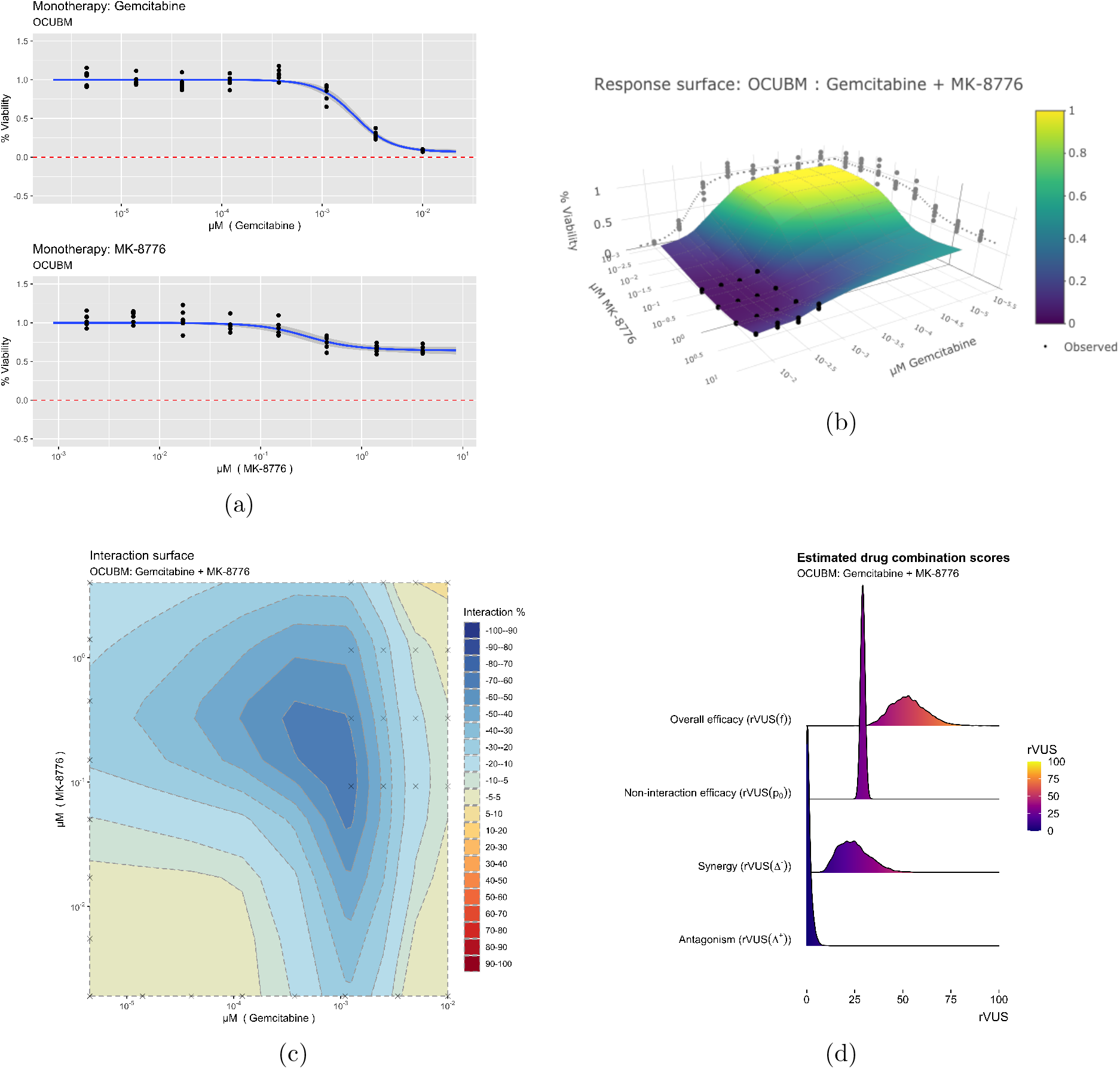
Some plots produced from the model output for the combination of Gemcitabine and MK-8776 on the OCUBM cell line. In panel (a), monotherapy curves for the two drugs, (b) a 3d interactive plot of the combined response, (c) contour plots of the interaction surface, and in (d) posterior densities of rVUS summaries.

For the overall efficacy, rVUS(*f*) is estimated at 53.38% of the maximum available volume, with a 95% CI (37.2,72.6). Of this effect, synergy (rVUS(Δ^−^)) accounts for about 25 percentage, points, 95% CI (11.8,43.5), while there is nearly no antagonistic effect at all (posterior mean of rVUS(Δ^+^) at 0.9). Note that the final estimate of synergy obtained by full MCMC sampling is larger than the initial estimate from the screen using the variational approximation. Variational inference algorithms are known for underestimating variances, which can explain why this effect was left underexplored. The wide credible interval in the synergy estimate is due to a large portion of the estimated effect being outside of the data range. Note that in Figure 5(c), the crosses denote the observed viability locations for the drug concentrations. The large synergistic effect is supported by 3 or 4 locations in the drug concentration landscape, with the bulk of the effect taking place outside of the data range. In this region, synergy is extrapolated using the underlying smoothness assumptions of the GP, as encoded by its length-scale *ℓ*. The variational approximation can underestimate the variability of this smoothness parameter, and thus also synergistic effects outside of the data range. We therefore recommend that the user rerun the most interesting experiments with full MCMC to better explore the uncertainties. It would also be natural to extend this experiment with slightly smaller concentrations of the drug combination, and in general there is little sense in looking for synergy in areas where the mono-therapies themselves are effective.

#### 5.1.1 Model assessment

The interaction part of the model consists of the GP, which is constructed from the latent parameters **z** together with the kernel hyperparameters 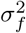 and *ℓ*, and the parameters inside the transformation, (*b*_1_, *b*_2_). Of these, only the kernel hyperparameters can be given a clear interpretation and are therefore reported in Table 5. We see that the length-scale parameter is estimated at 1.68, 95% CI (0.96,3.09), indicating a rather smooth function over the dose concentrations that range from 10^−6^*μ*M to 4*μ*M, also visible in Figure 5(c). The kernel amplitude 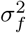 has a posterior mean of 3.69, with a wide 95% CI (1.32,13.35) indicating that the posterior function is allowed to deviate far from its mean, i.e. there is most likely interaction here as the function is allowed to deviate from *p*_0_. These parameters have a slightly smaller effective sample size, most likely due to the difficulty of updating these at the same time as updating **z**, as reported in [21]. While these parameters may not be of immediate interest to the user, they provide, together with the estimate of the observation noise *σ*, information about the model fit. Particularly when compared with other experiments across a large screen.

Having obtained estimates of synergy, which take the full uncertainty of the data into account, the user is left with a list of interesting experiments to follow-up. Because the model properly handles the estimation uncertainty, the list of such experiments is concise, focusing on those where a true effect can be clearly differentiated from the background noise. From here, the user can analyse the drug combination screen either qualitatively by going further into the biology, or quantitatively by plugging the output of the model into other algorithms for further analysis. One such avenue might be bio-marker discovery models as in the style of [28], who utilises both the efficacy estimate and its corresponding uncertainty to find bio-markers of single therapy responses. The output can also be used as training data for machine learning algorithms attempting to predict the combined drug efficacy or synergy for untested experiments.

## 6 Conclusion and outlook

The **bayesynergy** package implements a probabilistic model for analysing drug combination experiments. It handles the real world structure of drug combination experiments, that feature different patterns of missingness and differing numbers of replicates. The model accounts for measurement noise by using a heteroscedastic observation model, and produces estimates of the underlying dose-response function and measures derived from it, alongside uncertainty quantification. Since the model samples from the dose-response function directly, any user-defined summary statistic of dose-response can also be computed, and the uncertainty is naturally propagated.

Through the natural grid-structure of drug-combination experiments, a computationally efficient Gaussian Process implementation ensures the automatic exploration of the complete dose-response matrix from an incomplete input. This enables experimenters to analyse sparsely observed experiments in situations with limited resources, for example when using patient derived cell samples with limited biopsy material, to inform treatment decisions under uncertainty.

Cell viability assays are noisy by nature, with multiple sources of biological and technical noise. It is crucial to take this into account when estimating the efficacy and synergy of drug combinations in an attempt at understanding the underlying biological mechanisms. It enables researchers to hone in on the interesting combinations for follow-up experiments, and informs decision making and experimental design. More precise estimates of drug efficacy can be used as input in various models attempting to connect efficacy to underlying genomics patterns, or used to predict the response in new experiments or in patients.

In this paper, we have focused on the Bliss independence model as the underlying non-interaction assumption. This choice was motivated partly by the attractive probabilistic interpretation, but also by computational considerations. The Bliss model can be computed analytically from the two dose-response functions, while other models require numerical solutions (e.g. the Loewe model [1]). In an MCMC setting, this becomes expensive, as a numerical solver would need to be run at each step of the algorithm. Furthermore, both the NUTS algorithm and the variational approximation require the evaluation of the gradient of the log-posterior. Other models of non-interaction can introduce discontinuities in these gradients (e.g. the *Highest Single Agent* [29]), which would make posterior sampling slow and inefficient.

An advantage of the fully Bayesian model is that the user can explore various non-interaction assumptions *post-hoc*. From the posterior samples of the mono-therapy parameters, the user can construct various non-interaction assumptions to obtain 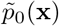, that can then be subtracted from the posterior dose response matrix in Figure 2 to obtain 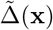. In this fashion, more suitable pharmacokinetic assumptions can be incorporated if desired.

Researchers working with drug efficacy screens frequently run multiple versions of their experiments. For example, in an initial phase of setting up a large drug screen, the active ranges of various drugs might need to be determined experimentally. Sometimes entire experiments are scrapped, because they fail to meet a quality control threshold. The **bayesynergy** model can readily be extended to not only consider within-experiment variability, but also between-experiment variability, as in [14]. This would allow the pooling of experimental data in estimating drug efficacy and synergy, utilizing all available data for the final analysis. The differences in experimental quality can be handled by assigning different weights to different experiments according to assay quality, as characterised by the positive and negative controls.

Finally, the extension of the model to higher orders of drug combinations is fairly straight forward, and still computationally efficient due to the grid structure. Considering 6 unique concentrations for each drug, a single-replicate fully observed drug combination experiment would contain 36, 216 and 1296(!) viability measurements for two, three, and four drugs respectively. In these settings resource constraints will quickly become an issue, and sparse designs an essential tool in the exploration of higher order drug synergies.

## 7 Key points

- Drug combination experiments are typically fraught with noise, both biological and technical.
- Accounting for this noise is key when searching for synergistic drug combinations in large screens, or when using estimates of synergy as input in other algorithms.
- The **bayesynergy** package implements a probabilistic model using Gaussian processes for analysing drug combination experiments, controlling for these noise sources.
- Since it is a statistical model, it allows inclusion of replicates, missing data and uneven concentration grids, in addition to providing uncertainty quantification around the results.

## 8 Software availability

The R package is available at https://github.com/ocbe-uio/bayesynergy. Scripts and datasets for reproducing Figures 4 and 5 and Table 5 can also be found there.

## 9 Appendix

## Detailed model specification

In the following, the full model is described in more details, including the specification of prior distributions for all the parameters.

Starting from the observation model, there are two parameters that need to be specified: the standard deviation *σ*, which is given an Inverse-Gamma(5,1) distribution, ensuring that its support includes reasonable levels of measurement error in proliferation assays, and the parameter *λ*, which is set to 0.005 by default, and can be tuned by the user to reflect the technical noise level in the assay.

The mono-therapy curves in the definition of *p*_0_ contain three parameters, slope (*s*), EC50 (*m*) and the lower asymptote (*l*). The slope parameter *s* is assumed to be positive, meaning that the dose-response function is monotonically decreasing with increasing drug concentration, the assumption being that no drugs are beneficial to cancer growth. This parameter is given a Gamma(1,1) prior, reflecting a preference for lower values of the slope (drug activation is smooth across the concentration range), but allowing for larger values and abrupt activation. The EC50 parameter *m* is a location parameter on the log_10_ scale, and is given a Normal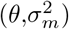 distribution, where 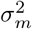 is modelled as Inverse-Gamma(3,2). To ensure a good fit, even in cases where the cell sample is unresponsive, we put a standard-normal distribution on *θ*. Finally, the lower asymptote *l* is given a Beta(1,1.25) prior, which places minimal weight on values close to one, allowing for a better identification of the slope and EC50 parameters. In the case where a compound is known to be cytotoxic, the user may also choose to fix this parameter to zero.

For the interaction term, we need to consider the parameters of the kernel function, as well as the two parameters (*b*_1_, *b*_2_) appearing in the transformation. The GP’s kernel function usually has two parameters, an amplitude 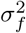 and a length-scale *ℓ* that controls the behaviour of the sample paths. By default, we utilize a Matèrn 3/2 kernel that takes the following form:

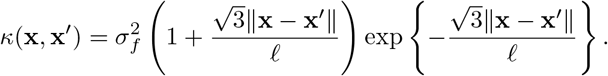

The amplitude 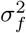 acts like a scale parameter on the GP sample paths, large values corresponding to function samples that can be far away from the mean function. Because the transformation function in equation (7) squeezes and centers the GP at *p*_0_, this parameter is given a heavy-tailed log-Normal(1,1) prior distribution allowing the model to express large synergistic or antagonistic values, when necessary. The length-scale *ℓ* controls the smoothness of the GP, with small values corresponding to rapid changes in output with minor changes in concentration. For this reason, *ℓ* is given an Inverse-Gamma(5,5) distribution penalizing values that are either too small or too large. Finally, the coefficients (*b*_1_, *b*_2_) in the GP transform are given independent Normal(1,0.1^2^) priors. This sharp prior ensures that the prior expectation of Δ(**x**) exists in a tight bound around zero (see Figure 6), and also avoids issues of non-identifiability together with 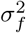.

**Figure 6:**
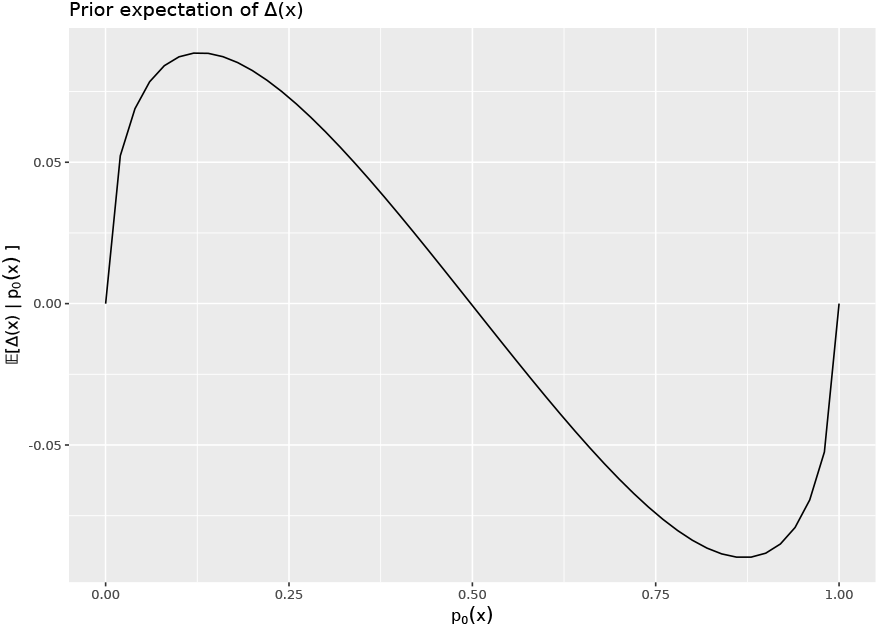
Figure showing the conditional prior expectation of Δ(**x**) given *p*_0_, as a function of *p*_0_(**x**)

Summarizing, we can write the full model specification as follows:

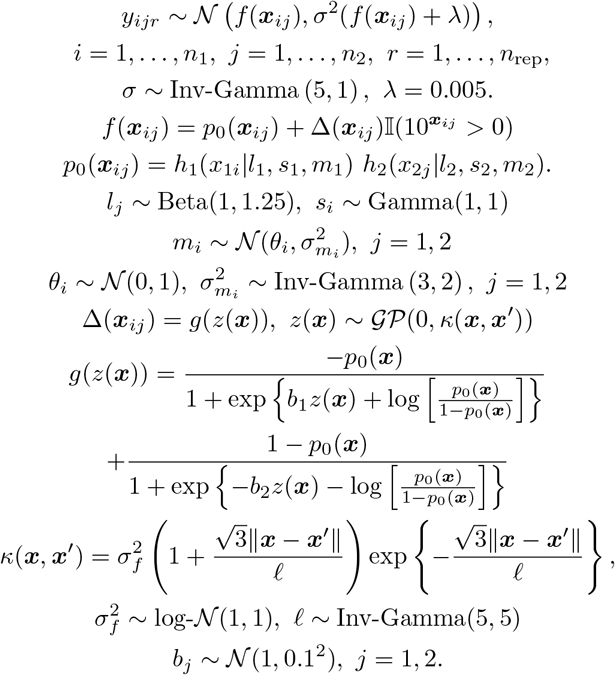

Note that the interaction term in the dose-response function, *f*, is multiplied with an indicator function, taking the value 1, if both concentrations are larger than zero on the original concentration scale and zero otherwise, ensuring that the model does not estimate synergy within the mono-therapy responses.

## Prior expectation of interaction

Figure 6 shows the prior expectation of Δ(**x**) conditional on values of *p*_0_(**x**) ranging from zero to one. The conditional expectation is calculated by sampling from the prior distribution, for fixed values of *p*_0_.

## References

[1] S. Loewe and H. Muischnek. Über Kombinationswirkungen. Naunyn-Schmiedebergs Archiv für experimentelle Pathologie und Pharmakologie, 114:313–326, 1926.

[2] C. I. Bliss. The toxicity of poisons applied jointly. Annals of applied biology, 26(3):585–615, 1939.

[3] W. R. Greco, G. Bravo, and J. C. Parsons. The search for synergy: a critical review from a response surface perspective. Pharmacological Reviews, 47(2):331–385, 1995.

[4] J. Fouquier and M. Guedj. Analysis of drug combinations: current methodological landscape. Pharmacology Research and Perspectives, 3(3), 2015.

[5] Christian T Meyer, David J Wooten, B Bishal Paudel, Joshua Bauer, Keisha N Hardeman, David Westover, Christine M Lovly, Leonard A Harris, Darren R Tyson, and Vito Quaranta. Quantifying drug combination synergy along potency and efficacy axes. Cell Systems, 8(2):97–108, 2019.

[6] Liye He, Evgeny Kulesskiy, Jani Saarela, Laura Turunen, Krister Wennerberg, Tero Aittokallio, and Jing Tang. Methods for High-Throughput Drug Combination Screening and Synergy Scoring, chapter 17, pages 351–398. Springer New York, 2018.

[7] Aleksandr Ianevski, Anil K Giri, and Tero Aittokallio. SynergyFinder 2.0: visual analytics of multi-drug combination synergies. Nucleic Acids Research, 48(W1):W488–W493, April 2020.

[8] C. Ritz, F. Baty, J. C. Streibig, and D. Gerhard. Dose-response analysis using R. PLOS ONE, 10(e0146021), 2015.

[9] Giovanni Y. Di Veroli, Chiara Fornari, Dennis Wang, Séverine Mollard, Jo L. Bramhall, Frances M. Richards, and Duncan I. Jodrell. Combenefit: an interactive platform for the analysis and visualization of drug combinations. Bioinformatics, 32(18):2866–2868, 04 2016.

[10] Arnaud Amzallag, Sridhar Ramaswamy, and Cyril H. Benes. Statistical assessment and visualization of synergies for large-scale sparse drug combination datasets. BMC Bioinformatics, 20(1), February 2019.

[11] Andrea Cremaschi, Arnoldo Frigessi, Kjetil Taskén, and Manuela Zucknick. A Bayesian approach for the study of synergistic interaction effects in *in-vitro* drug combination experiments. arXiv preprint arXiv:1904.04901, 2019.

[12] B. Yadav et al. Quantitative scoring of differential drug sensitivity for individually optimized anticancer therapies. Scientific reports, 4(5193), 2014.

[13] C. K. Williams and C. E. Rasmussen. Gaussian processes for machine learning. the MIT Press, 2(3):4, 2006.

[14] V. G. Hennessey, G. L. Rosner, R. C. Bast, and M. Y. Chen. A Bayesian approach to dose-response assessment and synergy and its application to in vitro dose-response studies. Biometrics, 66(4):1275–1283, Dec 2010.

[15] Wesley Tansey, Kathy Li, Haoran Zhang, Scott W Linderman, Raul Rabadan, David M Blei, and Chris H Wiggins. Dose–response modeling in high-throughput cancer drug screenings: an end-to-end approach. Biostatistics, January 2021.

[16] Ji-Hu Zhang, Thomas D. Y. Chung, and Kevin R. Oldenburg. A simple statistical parameter for use in evaluation and validation of high throughput screening assays. Journal of Biomolecular Screening, 4(2):67–73, April 1999.

[17] Bob Carpenter, Andrew Gelman, Matthew D. Hoffman, Daniel Lee, Ben Goodrich, Michael Betancourt, Marcus Brubaker, Jiqiang Guo, Peter Li, and Allen Riddell. Stan: A probabilistic programming language. Journal of Statistical Software, Articles, 76(1):1–32, 2017.

[18] Stan Development Team. RStan: the R interface to Stan, 2020. R package version 2.21.2.

[19] Matthew D. Hoffman and Andrew Gelman. The No-U-Turn Sampler: Adaptively setting path lengths in Hamiltonian Monte Carlo. Journal of Machine Learning Research, 15(47):1593–1623, 2014.

[20] Alp Kucukelbir, Rajesh Ranganath, Andrew Gelman, and David M. Blei. Automatic variational inference in Stan, 2015.

[21] Seth Flaxman, Andrew Gelman, Daniel Neill, Alex Smola, Aki Vehtari, and Andrew Gordon Wilson. Fast hierarchical Gaussian processes. 2015.

[22] Medhat Shehata, Susanne Schnabl, Dita Demirtas, Martin Hilgarth, Rainer Hubmann, Elena Ponath, Sigrun Badrnya, Claudia Lehner, Andrea Hoelbl, Markus Duechler, Alexander Gaiger, Christoph Zielinski, Josef D. Schwarzmeier, and Ulrich Jaeger. Reconstitution of PTEN activity by CK2 inhibitors and interference with the PI3-K/Akt cascade counteract the antiapoptotic effect of human stromal cells in chronic lymphocytic leukemia. Blood, 116(14):2513–2521, October 2010.

[23] Aleksandr Ianevski, Anil K. Giri, Prson Gautam, Alexander Kononov, Swapnil Potdar, Jani Saarela, Krister Wennerberg, and Tero Aittokallio. Prediction of drug combination effects with a minimal set of experiments. Nature Machine Intelligence, 1(12):568–577, December 2019.

[24] J. O’Neil et al. An unbiased oncology compound screen to identify novel combination strategies. Molecular Cancer Therapeutics, 15(6):1155–1162, 2016.

[25] Michael Choi, Thomas Kipps, and Razelle Kurzrock. ATM mutations in cancer: Therapeutic implications. Molecular Cancer Therapeutics, 15(8):1781–1791, July 2016.

[26] Adil I. Daud, Michelle T. Ashworth, Jonathan Strosberg, Jonathan W. Goldman, David Mendelson, Gregory Springett, Alan P. Venook, Sabine Loechner, Lee S. Rosen, Frances Shanahan, David Parry, Stuart Shumway, Jennifer A. Grabowsky, Tomoko Freshwater, Christopher Sorge, Soonmo Peter Kang, Randi Isaacs, and Pamela N. Munster. Phase I dose-escalation trial of checkpoint kinase 1 inhibitor MK-8776 as monotherapy and in combination with gemcitabine in patients with advanced solid tumors. Journal of Clinical Oncology, 33(9):1060–1066, March 2015.

[27] S Origanti, S r Cai, A Z Munir, L S White, and H Piwnica-Worms. Synthetic lethality of Chk1 inhibition combined with p53 and/or p21 loss during a DNA damage response in normal and tumor cells. Oncogene, 32(5):577–588, March 2012.

[28] Dennis Wang, James Hensman, Ginte Kutkaite, Tzen S Toh, Ana Galhoz, Howard Lightfoot, Wanjuan Yang, Maryam Soleimani, Syd Barthorpe, Tatiana Mironenko, Alexandra Beck, Laura Richardson, Ermira Lleshi, James Hall, Charlotte Tolley, William Barendt, Jonathan R Dry, Julio Saez-Rodriguez, Mathew J Garnett, Michael P Menden, and Frank Dondelinger and. A statistical framework for assessing pharmacological responses and biomarkers using uncertainty estimates. eLife, 9, December 2020.

[29] Joseph Lehár, Grant R Zimmermann, Andrew S Krueger, Raymond A Molnar, Jebediah T Ledell, Adrian M Heilbut, Glenn F Short, Leanne C Giusti, Garry P Nolan, Omar A Magid, Margaret S Lee, Alexis A Borisy, Brent R Stockwell, and Curtis T Keith. Chemical combination effects predict connectivity in biological systems. Molecular Systems Biology, 3(1):80, January 2007.

